# MucOneUp: A Simulation Framework for *MUC1*-VNTR Variant Benchmarking

**DOI:** 10.64898/2026.05.08.723876

**Authors:** Bernt Popp, Hassan Saei

## Abstract

**Summary:** Variable number tandem repeats (VNTRs) in the *MUC1* gene cause autosomal dominant tubulointerstitial kidney disease when disrupted by frameshift variants, but the GC-rich 60-bp repeat structure (20-125 copies) challenges variant detection. While tools like VNtyper enable *MUC1* variant calling, no gold-standard benchmarking datasets exist for systematic performance evaluation. We present MucOneUp, a specialized simulation framework for generating *MUC1*-VNTR reference sequences with targeted variants and platform-specific sequencing reads (Illumina, Oxford Nanopore, PacBio). MucOneUp employs Markov chain-based repeat generation, supports diploid simulation with customizable variant placement, and includes additional analysis modules for SNaPshot assay simulation and exploratory frameshift analysis. We validate MucOneUp through a multi-variant, cross-platform benchmark of six tool-platform combinations using 13 distinct frameshift variants and investigate VNTR length effects on detection.

**Availability and implementation:** MucOneUp is accessible at no cost under the MIT License at https://github.com/berntpopp/MucOneUp and archived on Zenodo (DOI: 10.5281/zenodo.19740406).

**Supplementary information:** Supplementary data are provided with this manuscript.

## Introduction

The *MUC1* variable number tandem repeat (VNTR) consists of a 60-bp GC-rich unit repeated 20-125 times, encoding the MUC1 protein extracellular domain (Kirby *et al*., 2013; Logsdon *et al*., 2020). Frameshift variants in this repeat cause autosomal dominant tubulointerstitial kidney disease (ADTKD-*MUC1*), producing progressive renal fibrosis and early-onset chronic kidney disease (Kirby *et al*., 2013; Ekici *et al*., 2014).

The resulting truncated proteins are trapped in TMED9-containing vesicles of the early secretory pathway and trigger cellular toxicity (Dvela-Levitt *et al*., 2019).

Clinical diagnosis often depends on targeted assays for the recurrent 59dupC cytosine duplication, historically also reported as 27dupC or c.428dupC and implemented in MucOneUp as dupC (Ekici *et al*., 2014). The SNaPshot minisequencing assay, for example, requires multiple digestion and PCR steps, is prone to contamination, and few laboratories offer it; it also cannot detect novel frameshift events outside the targeted region (Ekici *et al*., 2014). Long-read sequencing can now target the full-length VNTR using primers first described by Kirby et al. (Kirby *et al*., 2013) and later adopted in long-range amplicon workflows (Wenzel *et al*., 2018; Vrbacká *et al*., 2026; Madritsch *et al*., 2025). However, reusable analysis code and standardized validation datasets remain limited, complicating routine clinical benchmarking (Kachmar *et al*., 2025).

Short-read computational methods such as VNtyper (Saei *et al*., 2023) and codeadVNTR (Park *et al*., 2022) detect *MUC1* variants using k-mer-based alignment-free methods (Audano *et al*., 2018) or hidden Markov models (Bakhtiari *et al*., 2018), allowing screening from standard diagnostic sequencing. However, unlike variant types covered by community benchmarks such as the Genome in a Bottle consortium (Zook *et al*., 2019), no gold-standard datasets with known ground truth exist for *MUC1*-VNTR, preventing systematic performance assessment across platforms.

Simulation could provide synthetic ground truth, but existing read simulators (Wessim2 (Kim *et al*., 2013; Wang and Kong, 2019), NanoSim (Yang *et al*., 2017), PBSIM3 (Ono *et al*., 2022)) use haploid reference sequences and cannot model diploid haplotypes with different VNTR lengths or place variants at specific repeat positions. No existing tool provides exploratory frameshift analysis or SNaPshot assay accessibility assessment for this locus.

To provide locus-specific ground truth for *MUC1*-VNTR method evaluation, we developed MucOneUp. It generates diploid reference sequences with variable VNTR structures using Markov chain transitions, places specific variants at defined repeat positions, simulates sequencing reads with established platform-specific simulators, and includes modules for SNaPshot assay simulation and exploratory frameshift analysis. Here we describe its implementation and validate it through cross-platform benchmarking of six tool-platform combinations.

## Methods

We implemented MucOneUp in Python as a modular pipeline that generates diploid haplotypes, inserts targeted variants, and simulates reads across multiple sequencing platforms. Additional modules provide SNaPshot assay accessibility assessment and exploratory frameshift analysis. Users configure simulations through JSON Schema-validated files that specify repeat definitions, Markov transition probabilities, variant catalogs, and platform parameters (Supplementary Methods SM1).

### VNTR Generation

MucOneUp generates VNTR structures using discrete-time Markov chain transitions (Durbin *et al*., 1998). Each transition depends only on the current repeat unit (first-order Markov process). The *MUC1* configuration employs 34 distinct repeat units curated from the published literature and following the naming convention introduced in the initial description (Kirby *et al*., 2013). These 34 repeat units were calibrated to frequencies from published repeat compositions (Wenzel *et al*., 2018). All haplotypes enforce a canonical terminal block (6 or 6p → 7 → 8 → 9) that matches the conserved C-terminal structure of wildtype *MUC1* alleles. The variable region is dominated by repeat unit X (transition probability ∼0.7489), with minor contributions from A, B, C, D, and other letter repeats. Weighted transitions produce sequences with biological composition and natural variability.

### Diploid Simulation

The tool generates independent haplotypes with configurable length parameters (normal distribution with bounded range, or fixed lengths via CLI) using a random seed for reproducibility. Reference sequences contain flanking regions (10 kb each) and the VNTR array. The tool supports hg19 and hg38 coordinates. FASTA headers encode haplotype identity and variant annotations; full metadata is recorded in the companion simulation_stats.json file (Supplementary Methods SM1).

### Variant Application

Variants are applied using 1-based repeat indexing with validation against allowed repeat types. Dual-mode simulation produces matched wildtype and variant-containing outputs from the same simulated haplotypes, ensuring identical VNTR structures for comparative benchmarking. Strict mode ensures variant placement fidelity by rejecting invalid targets. Each simulation records full metadata (SHA-256 configuration fingerprint, random seeds, haplotype lengths, variants, software version) for reproducibility (Supplementary Methods SM1).

### Read Simulation

Integration with established simulators provides platform-specific error profiles. Illumina reads are simulated using Wessim2-style fragment generation (Kim *et al*., 2013; Wang and Kong, 2019) with ReSeq2 error modeling (a maintained fork of ReSeq (Schmeing and Robinson, 2021)). PBSIM3 (Ono *et al*., 2022) simulates PacBio HiFi and ONT amplicon reads; NanoSim (Yang *et al*., 2017) is integrated for whole-genome ONT simulation. Both amplicon pipelines simulate haplotypes independently: coverage is split between alleles according to a PCR length-bias model (favouring the shorter haplotype), each haplotype is simulated with a unique seed, and reads are merged. Whole-genome ONT simulation follows a separate regime: NanoSim’s length-dependent read generation is wrapped in a diploid split-simulation that divides target coverage equally between haplotypes, reducing naive allelic bias from 3.27:1 to 1.40:1. All pipelines generate aligned and indexed BAM files using BWA (Li and Durbin, 2009) (Illumina) or minimap2 (Li, 2018) (Oxford Nanopore and PacBio) and SAMtools (Li *et al*., 2009) (Supplementary Methods SM1). For the benchmark, we used target depths of 150x for Illumina short reads and 500x for PacBio/ONT amplicon reads; these experiment-level settings are summarized in Supplementary Methods SM2 and can be changed through the MucOneUp configuration.

### Additional Analysis Modules

Two modules analyze properties of simulated variants. The SNaPshot assay validator simulates the minisequencing protocol (Ekici *et al*., 2014) through *in silico* PCR with multi-product handling, MwoI restriction digest, and single-base extension. This predicts whether variants at different VNTR positions are detectable by the current clinical assay. The exploratory frameshift analyzer scores ORF-predicted protein sequences by combining repeat pattern detection with amino acid composition analysis (Figure 1 B). The composite score uses heuristic weights (repeat 0.6, composition 0.4, threshold 0.5) chosen to prioritize repeat structure; these are not empirically optimized. This module generates hypotheses about frameshift consequences and lacks clinical validation (Supplementary Methods SM1).

**Figure 1:**
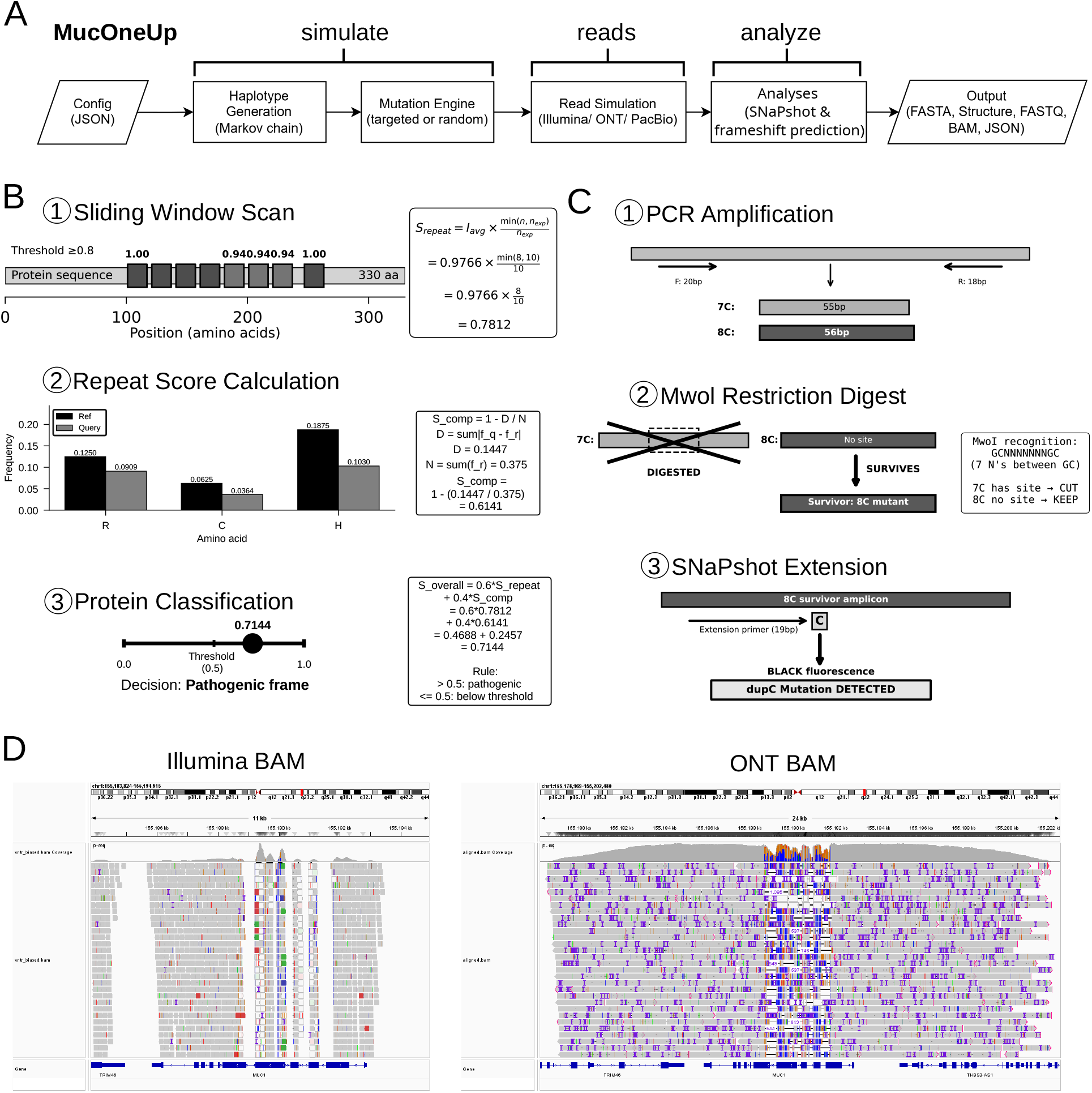
MucOneUp workflow and validation. **A** CLI subcommands (simulate, reads, analyze) with JSON inputs and outputs (FASTA, BAM, predictions). **B** Exploratory frameshift analysis via (1) sliding window repeat detection, (2) repeat score calculation, and (3) composite score computation. **C** SNaPshot simulation: (1) PCR amplification, (2) MwoI restriction digest, and (3) SNaPshot extension with variant detection. **D** IGV screenshots of simulated BAM files: Illumina exome and Oxford Nanopore reads showing platform-specific coverage.

## Results

### *MUC1* VNTR Variant Caller Benchmarking

We benchmarked *MUC1* VNTR variant calling on 220 MucOneUp-generated pairs carrying 13 frameshift variants (100 59dupC pairs and 10 pairs each for 12 atypical variants) with mean 50 repeats per haplotype (Supplementary Figure S1, Methods SM2, SM4). We evaluated six tool-platform combinations across Illumina short reads (150x) and PacBio/ONT amplicon long reads (500x). Table 1 summarizes the results. VNtyper 2 Fast and Normal modes performed identically; VNTRPipeline (Madritsch *et al*., 2025) achieved the highest overall sensitivity via assembly-based LOF detection on ONT reads. Per-variant sensitivity ranged from 30% (insA_pos54) to 100% (insCCCC, insG_pos54, insG_pos58) for VNtyper 2 Fast, with 59dupC at 93% (Supplementary Figure S1).

**Table 1.** Cross-platform benchmarking (220 pairs, 13 variants). Normal omitted (=Fast). Sensitivity and Specificity CIs = Wilson 95%; F1 CI = 10,000-resample bootstrap 95%.

| Tool (Platform) | Sens. [95% CI] | Spec. [95% CI] | F1 [95% CI] |
| --- | --- | --- | --- |
| VNtyper 2 Fast (Illumina) | 0.8727 [0.8222, 0.9105] | 1 [0.9828, 1.0000] | 0.932 [0.9051, 0.9556] |
| VNtyper 2 Shark (Illumina) | 0.8818 [0.8325, 0.9181] | 1 [0.9828, 1.0000] | 0.9372 [0.9123, 0.9600] |
| VNtyper 2 adVNTR (Illumina) | 0.5955 [0.5295, 0.6581] | 1 [0.9828, 1.0000] | 0.7464 [0.6925, 0.7961] |
| open-pacmuci (PacBio) | 0.8409 [0.7868, 0.8833] | 0.9455 [0.9071, 0.9685] | 0.8873 [0.8526, 0.9180] |
| open-pacmuci (ONT) | 0.5091 [0.4434, 0.5744] | 1 [0.9828, 1.0000] | 0.6747 [0.6129, 0.7313] |
| VNTRPipeline (ONT) | 0.9495 [0.9119, 0.9716] | 0.9908 [0.9672, 0.9975] | 0.9696 [0.9515, 0.9845] |

False negatives (n=28 for Fast/Normal, n=26 for Shark, n=89 for adVNTR) concentrated among atypical variants and samples with long alleles (longest allele 52-150 repeats for Kestrel modes, 33-150 for adVNTR; Figure S1). All Kestrel false negatives had zero depth score, while all true positives scored above 4.7e-3.

### Paired Enrichment Analysis

We evaluated the VNtyper 2 count-based implementation of the Mutation Counter method (Fages *et al*., 2024) on the same simulated Illumina pairs. The module counts fixed target sequences and their reverse complements directly from BAM reads before Fisher testing against the All count column (Supplementary Figure S2, Methods SM3). The predefined panel of variants covered 140/220 benchmark pairs; the remaining 80 pairs carried variants outside that panel. Among evaluable pairs, 138/140 (98.6%) showed significant enrichment at *P*<1.0e-3.

### Detection Sensitivity Across VNTR Lengths

We analyzed 72 diploid 59dupC samples (3 replicates per length) with fixed Haplotype 1 (60 repeats) and variable Haplotype 2 (20-130 repeats, 10-repeat increments). The 59dupC variant was placed on Hap1 (Condition A, n=36) or Hap2 (Condition B, n=36) (Supplementary Figure S3, Methods SM2). VNtyper 2 Fast mode detected 68/72 (94%) variants, with missed calls at the longest VNTR lengths.

Both conditions showed strong negative correlations between depth score and variable allele length (Condition A: Pearson r=-0.693, *P*=2.86 × 10^−6^; Condition B: Pearson r=-0.625, *P*=4.54 × 10^−5^; Supplementary Methods SM4). Fisher’s Z-test showed no significant difference between conditions (z=-0.49, *P*=0.6244), indicating that allele length affects detection regardless of which haplotype carries the variant. Allele length is therefore the dominant factor in missed calls, not variant placement.

### Systematic Frameshift Analysis

We applied MucOneUp’s ORF analysis to 30 synthetic variants (insertions and deletions of 1–15 bp) at the canonical 59dupC position. All 10 in-frame controls (multiples of 3 bp) scored below the composite threshold; of 20 out-of-frame variants, 10 in the pathogenic frame were classified as pathogenic-frame and 10 in the non-pathogenic frame scored below threshold (Supplementary Table S5). The 59dupC-equivalent one-cytosine insertion, labelled `ins1C` in this synthetic ORF series, produced a 88 AA truncation with composite score 0.82.

## Discussion

Our benchmarking reveals that *MUC1* variant detection performance varies across tools and platforms, with VNTR allele length as the dominant confounder. False negatives concentrated among atypical variants and long alleles, consistent with the difficulty of resolving k-mer signals in extended repetitive sequences. *MUC1* frameshifts produce two reading frames: one generates a truncated protein (∼85 amino acids downstream) that presumably accumulates in the early secretory pathway (Dvela-Levitt *et al*., 2019), while the other terminates later (∼189 amino acids) and is presumably degraded (Supplementary Table S4). The cellular fate of these frame-specific products remains incompletely defined, so we interpret the ORF scores as exploratory sequence-based prioritization rather than toxicity predictions. In the synthetic frameshift series, all in-frame controls scored below threshold, whereas pathogenic-frame out-of-frame variants were prioritized (Supplementary Table S5).

Cross-platform comparison shows that assembly-based long-read approaches like VNTRPipeline (Madritsch *et al*., 2025) compensate for ONT error rates, while alignment-based tools show platform-dependent sensitivity. To avoid simulation artefacts, MucOneUp separates haplotypes, balances coverage, and simulates independently rather than sampling concatenated references with length-proportional probability. The long-read tools evaluated here all rely on the same amplicon protocol, yet none shipped with shared validation data. We reimplemented Vrbacka’s PacBio workflow from the published description (Vrbacká *et al*., 2026) as open-pacmuci and validated it using MucOneUp synthetic data. MucOneUp also simulates amplicons using the same primers, providing benchmarking capability for a protocol that has lacked standardized validation since its initial description.

We calibrated simulated haplotypes to published Wenzel repeat-unit compositions (Figure S9; Table S7) (Wenzel *et al*., 2018) and anchored benchmark lengths to the Vrbacka PacBio distribution (Methods SM2) (Vrbacká *et al*., 2026). Because no orthogonally curated diagnostic haplotypes are available, these checks support source-data realism rather than independent biological validation. Transition probabilities derive from one predominantly European-descent study (Wenzel *et al*., 2018) and may not capture full population diversity; alternative locus-specific matrices can be regenerated from user-supplied composition data via the `analyze vntr-stats` subcommand. For Illumina, the tool models VNTR capture efficiency using an empirically derived penalty but not panel-specific probe biases beyond our Twist v2 profile. For amplicon sequencing, MucOneUp models length-dependent PCR amplification bias but not primer dimers, PCR chimeras, or off-target amplification.

While calibrated for *MUC1*, the Markov chain framework could be adapted to other VNTRs with heterogeneous repeat composition using locus-specific transition matrices. As long-read sequencing tools for *MUC1* continue to develop, MucOneUp provides standardized benchmarking datasets to evaluate their analytical performance; MucOneUp supports background SNP injection for polymorphism-aware simulations (Supplementary Methods SM1).

## Availability and Implementation

MucOneUp is implemented in Python 3.10+ and available under MIT License at https://github.com/berntpopp/MucOneUp, archived on Zenodo (DOI: 10.5281/zenodo.19740406). Dependencies are managed via uv package manager; external tools (ReSeq2, NanoSim, PBSIM3, BWA (Li and Durbin, 2009), minimap2, SAMtools) are installable via conda or through the provided Docker image (ghcr.io/berntpopp/muconeup/muconeup). The tool includes a database of 13 known *MUC1* variants with literature citations, pre-configured repeat unit catalogs, and documentation with tutorial workflows. All outputs will be maintained for two years post-publication.

## Acknowledgments

We thank the open-source bioinformatics community for developing the simulation, error-modeling, alignment, consensus, and file-processing tools that make MucOneUp possible, including Wessim/Wessim2-style approaches, ReSeq2, NanoSim, PBSIM3, BWA-MEM, minimap2, CCS, and SAMtools.

## Author Contributions

Bernt Popp (Conceptualization [lead], Software [lead], Formal analysis [lead], Investigation [lead], Methodology [lead], Visualization [lead], Writing–original draft [lead], Writing–review & editing [equal]). Hassan Saei (Conceptualization [equal], Methodology [supporting], Writing–review & editing [equal]).

## Supplementary Data

Supplementary data are provided with this manuscript.

## Conflict of Interest

B.P. is employed by Labor Berlin – Charité Vivantes GmbH, a diagnostic laboratory offering genetic testing services. H.S. developed the original VNtyper implementation;

B.P. and H.S. developed VNtyper 2, which is benchmarked in this study. No other conflicts declared.

## Funding

None declared.

## Data Availability

The source code and all data underlying this Application Note are available at https://github.com/berntpopp/MucOneUp. Benchmarking data are deposited on Zenodo as two linked records: simulation inputs and simulated reads (DOI: 10.5281/zenodo.19741156), and tool outputs with analysis results (DOI: 10.5281/zenodo.19741672). Complete simulation parameters and benchmarking results are provided in Data S1. The open-pacmuci reimplementation is available at https://github.com/berntpopp/open-pacmuci. ReSeq2, the maintained fork of ReSeq used for Illumina error modeling, is available at https://github.com/berntpopp/ReSeq2.

## Declaration of Generative AI in Manuscript Preparation (46 words)

During the preparation of this work the authors used Anthropic Claude (Opus 4, Sonnet 4) and GitHub Copilot to assist with software development, data analysis scripting, figure generation, and language refinement. The authors reviewed and edited all content and take full responsibility for the published article.

## Notes

https://github.com/berntpopp/MucOneUp

https://github.com/berntpopp/open-pacmuci

https://github.com/berntpopp/ReSeq2

https://doi.org/10.5281/zenodo.19740406

https://doi.org/10.5281/zenodo.19741156

https://doi.org/10.5281/zenodo.19741672

https://doi.org/10.5281/zenodo.19744166

https://ghcr.io/berntpopp/muconeup/muconeup

### Competing Interest Statement

B.P. is employed by Labor Berlin - Charite Vivantes GmbH, a diagnostic laboratory offering genetic testing services. H.S. developed the original VNtyper implementation; B.P. and H.S. developed VNtyper 2, which is benchmarked in this study. No other conflicts are declared by any author.

