## Supplementary Materials for "MucOneUp: A Simulation Framework for *MUC1*-VNTR Variant Benchmarking"

### Supplementary Materials: MucOneUp - A Simulation Tool for *MUC1*-VNTR Variant Benchmarking and Diagnostic Validation

**2 Supplementary Figures** **18**

**3 Supplementary Tables** **25**

**4 Supplementary Data** **33**

**5 References** **34**

### 1 Supplementary Methods

#### 1.1 SM1: MucOneUp software implementation and simulation model

##### 1.1.1 Software architecture

MucOneUp (v0.44.4) comprises a simulation engine for Markov-chain VNTR construction, a variant engine that applies targeted edits to repeat units, multi-platform read simulators wrapping ReSeq2 (Illumina), NanoSim (whole-genome ONT), and PBSIM3 (PacBio HiFi and ONT amplicon), an ORF-translation and frameshift-scoring module, a SNaPshot-assay validator, and a reproducibility/provenance layer that records configuration fingerprints, seeds, and sanitized command-line provenance for every run.

**Installation:** `git clone https://github.com/berntpopp/MucOneUp.git && cd MucOneUp && make install`

**Documentation:** https://berntpopp.github.io/MucOneUp/

##### 1.1.2 Command-line workflow

MucOneUp follows Unix philosophy with strict command separation: `simulate` generates haplotype FASTA files, `reads` simulates sequencing reads, and `analyze` performs downstream analyses (ORF prediction, VNTR statistics, SNaPshot validation).

**Basic workflow:**

```
# Step 1: Generate haplotypes
muconeup --config config.json simulate \
  --out-base simulations/pair_001/sample \
  --fixed-lengths 60 --fixed-lengths 80 \
  --mutation-name dupC \
  --mutation-targets 1,25 \
  --seed 1000

# Step 2: Simulate reads (Illumina)
muconeup --config config.json reads illumina \
  simulations/pair_001/sample.001.simulated.fa \
  --coverage 150
```

**Dual-mode simulation** (wildtype + variant-containing haplotypes with synchronized seeds):

```
# Step 1: Generate both wildtype and variant-containing haplotypes
muconeup --config config.json simulate \
  --out-base simulations/pair_001/sample \
  --fixed-lengths 60 --fixed-lengths 80 \
  --mutation-name normal,dupC \
  --mutation-targets 1,25 \
  --seed 1000

# Produces: sample.001.normal.simulated.fa (wildtype)
```

```
#           sample.001.mut.simulated.fa    (variant-containing)

# Step 2: Simulate reads for both (separate commands)
muconeup --config config.json reads illumina \
  simulations/pair_001/sample.001.normal.simulated.fa \
  --coverage 150

muconeup --config config.json reads illumina \
  simulations/pair_001/sample.001.mut.simulated.fa \
  --coverage 150
```

###### 1.1.3 Reproducibility metadata

Each simulation generates a simulation_stats.json file combining haplotype-level statistics (repeat counts, VNTR lengths, GC content, variants) from the `simulation_statistics` module with provenance metadata (software version, SHA-256 configuration fingerprint, random seeds, timestamps, and sanitized command-line invocation with secrets redacted) from the `provenance` module.

###### 1.1.4 Core simulation model

MucOneUp represents a VNTR allele as an ordered chain of repeat-unit symbols and then assembles the DNA sequence by concatenating the left constant flank, repeat-unit sequences, and the right constant flank. The default configuration contains locus-specific repeat definitions, hg19/hg38 flanking constants, a length model, and a transition-probability matrix. For each haplotype, `simulate_diploid()` either uses user-specified fixed lengths or samples a repeat count from the configured bounded normal distribution (default range 20-130 repeats in the benchmark configuration). A seed initializes the random-number generator, allowing exact regeneration of the same haplotypes.

Repeat chains are generated with a first-order Markov model: at step *t*, the next repeat symbol is sampled from the probability row for the current symbol only. The transition matrix was derived from published *MUC1*-VNTR repeat compositions (Wenzel *et al.*, 2018) by counting observed adjacent repeat pairs and normalizing each source-symbol row. The `analyze vntr-stats` command can regenerate these probabilities from user-supplied VNTR structure tables. During construction of the variable region, terminal symbols (`6`, `6p`, `9`, and `END`) are filtered out so that the chain does not terminate prematurely. After the variable segment is built, MucOneUp appends the conserved terminal structure (6 or 6p -> 7 -> 8 -> 9), matching the canonical C-terminal block of wildtype *MUC1* alleles. The assembled haplotypes therefore combine stochastic internal repeat variation with a constrained terminal architecture.

Each diploid simulation produces two independently sampled haplotypes unless fixed chains or fixed lengths are supplied. The final FASTA contains haplotype-specific records, and the companion metadata records repeat counts, VNTR lengths, GC content, repeat chains, variant annotations, genomic coordinates, software version, configuration fingerprint, seed values, and sanitized command-line provenance.

**Markov-order limitation.** MucOneUp's first-order chain captures marginal repeat-to-repeat transition frequencies but not higher-order dependencies such as long-range block duplications or multimeric motifs that are known features of complex VNTRs.

##### 1.1.5 Variant model

Variants are defined in the JSON configuration as named records with allowed repeat symbols, one or more edit operations, optional literature metadata, and a strictness flag. `apply_mutations()` receives 1-based targets of the form `(haplotype_index, repeat_index)`, validates that the requested repeat exists, and checks whether the target repeat symbol is permitted for the selected variant. In strict mode, disallowed repeat targets raise an error. In permissive mode, the target repeat can be converted to a randomly selected allowed repeat symbol before applying the edit, which is useful for exploratory variant sweeps but less appropriate for locked benchmarking designs.

Within the selected 60-bp repeat unit, MucOneUp supports four edit classes: insertion, deletion, replacement, and deletion-insertion. Coordinates are defined relative to the repeat unit rather than the whole haplotype, which makes variants portable across alleles of different VNTR length. After editing, the affected repeat unit is marked as variant-bearing in the repeat chain (for example, `X` becomes `Xm` in structure outputs), the haplotype sequence is reassembled from the updated chain, and the exact target and altered repeat sequence are written to the metadata. Dual-mode simulation (`normal,<mutation>`) generates matched wildtype and variant-containing outputs from the same underlying haplotype structures, enabling paired benchmarking in which the only intended difference is the configured variant.

##### 1.1.6 Read simulation model

The `reads` command dispatches to platform-specific backends through a shared strategy-pattern interface, allowing Illumina, whole-genome ONT, PacBio, and amplicon-specific workflows to share a consistent CLI and output convention while keeping simulatorspecific logic isolated.

For Illumina data, MucOneUp uses a Wessim2-style fragment simulator implemented in Python rather than calling the original Wessim2 binary. The pipeline prepares the simulated FASTA, maps candidate fragments against the reference context, samples fragments according to the configured insert-size distribution (250 +/- 35 bp), and passes fragment templates to ReSeq2 for empirical Illumina error modeling using the Hs-Nova-TruSeq.reseq profile. ReSeq2 is a maintained fork of ReSeq (Schmeing and Robinson, 2021) and preserves the `.reseq` profile format. The resulting paired-end reads are aligned with BWA-MEM v0.7.17, sorted and indexed with SAMtools v1.21, and optionally downsampled. For exome-style simulations, MucOneUp uses a Twist Bioscience Exome v2 empirical capture profile plus a configurable VNTR capture-efficiency penalty (factor 0.39). This penalty is platform-specific; users simulating other exome kits or targeted panels should recalibrate it against their own coverage data.

For long-read amplicon data, MucOneUp first extracts per-haplotype PCR products using the configured primers from Wenzel *et al.* (2018) (forward: GGAGAAAAGGA-GACTTCGGCTACCCAG; reverse: GCCGTTGTGCACCAGAGTAGAAGCTGA; expected product range 500-15,000 bp). Diploid inputs are split into haplotype FASTAs, amplicons are extracted independently, and the total requested coverage is divided between alleles using the PCR length-bias model. The default deterministic model computes per-cycle amplification efficiencies as $E(L) = E_{\text{max}} \exp(-\alpha L)$, converts them into allele yields $(1 + E(L))^{cycles}$, and allocates read counts in proportion to those yields; this preferentially amplifies shorter amplicons while preserving a defined total coverage. Per-allele template FASTAs are then simulated independently with allele-specific seeds and merged before alignment.

The ONT amplicon workflow uses PBSIM3 v3.0.1 in template mode with the QSHMM-ONT-HQ model and minimap2 v2.24 `map-ont` alignment. The PacBio amplicon workflow uses PBSIM3 with the ERRHMM-SEQUEL model, generates multi-pass reads, applies CCS v6.4.0 consensus generation, and aligns HiFi reads with minimap2 `map-hifi`. Separately, MucOneUp provides a whole-genome ONT mode built around NanoSim (Yang *et al.*, 2017). This mode was not used in the benchmark experiments; it applies a diploid split-simulation wrapper that simulates haplotypes separately with adjusted coverage to reduce length-proportional allelic sampling bias. Across platforms, the BAM outputs shown in the main manuscript figure are not merely illustrative: they are the alignment products used for downstream benchmarking and quality inspection.

###### 1.1.7 Auxiliary analysis modules

The `analyze` command includes two optional modules that are methodologically separate from read generation. This subsection defines their algorithms; experiment-specific outputs and sensitivity analyses are reported later in the supplementary figures and tables.

The exploratory frameshift analyzer predicts ORFs from simulated FASTA files and scores each candidate protein with the frameshift-scoring module. The score combines two normalized sequence features: (i) a repeat score, $S_{repeat} = I_{avg} \times \min(n, n_{expected})/n_{expected}$, where $I_{avg}$ is the mean identity of repeat-like windows to the 16 amino-acid MUC1 consensus (`RCHLGPGHQAGPGLHR`), $n$ is the number of detected windows, and $n_{expected}$ caps the count contribution; and (ii) an amino-acid composition score, $S_{composition} = 1 - \sum |f_q - f_r| / \sum f_r$, comparing query and reference frequencies for R, C, and H. The final score is $S_{overall} = w_r S_{repeat} + w_c S_{composition}$.

Scores at or above the configured threshold are flagged as pathogenic-frame candidates. This is a sequence-level prioritization heuristic for simulated frameshifts, not a clinically validated toxicity assay.

Sensitivity of the binary classification to score weights and thresholds is reported in Figure S7 and Table S6.

The SNaPshot validator models the canonical 59dupC/8C minisequencing assay (Ekici *et al.*, 2014) as a three-step decision chain. `PCRSimulator` enumerates primer-derived amplicons and applies size and variant-pattern filters. `DigestSimulator` applies the configured MwoI digest and retains only products lacking the recognition site. `SnapshotExtensionSimulator` performs single-base extension on surviving products and maps the incorporated base to the configured fluorescence channel. Detection is called when the observed extension signal matches the expected variant channel.

##### 1.1.8 Background SNP injection

MucOneUp supports background SNP injection via the `simulate` command. SNPs are supplied either from a TSV input (`--snp-input-file`) or generated at random (`--random-snps --random-snp-density <per-kb>`), with independent control over targeted region (`--random-snp-region {all|constants_only|vntr_only}`) and haplotype (`--random-snp-haplotypes {all|1|2}`). The module reports applied substitutions and reference-base mismatches.

Three use cases motivate the capability: (i) polymorphism-aware variant-caller benchmarking across the VNTR and flanks; (ii) MwoI-site disruption testing for the SNaPshot assay, in which constants-only SNPs are placed within or adjacent to MwoI recognition sequences to assess how site-disrupting polymorphisms degrade SNaPshot sensitivity, a known clinical failure mode; and (iii) de novo assembly evaluation under realistic background polymorphism. The benchmarks reported here (Experiments 1-3) do not exercise background polymorphism; haplotypes are simulated without SNP injection.

###### 1.1.9 VNTR composition calibration against published repeat-unit data

MucOneUp repeat-unit transitions were parameterized from the long-read-resolved *MUC1* VNTR alleles reported by Wenzel *et al.* (2018). We therefore compared simulated normal haplotypes with the Wenzel allele set to assess whether the simulator preserves the published repeat-unit composition distribution after length sampling, forbidden terminal-state filtering, and addition of the canonical terminal block. This analysis evaluates calibration to the source composition data rather than independent external validation.

For each allele, repeat-unit composition was represented as a 28-category proportion vector. Mean composition concordance was assessed using Pearson and Spearman correlation, total variation distance, Jensen-Shannon divergence, and Hellinger distance. Per-unit differences were tested using Mann-Whitney U tests with Benjamini-Hochberg correction. Multivariate structure was evaluated using PERMANOVA and PERMDISP on Bray-Curtis distances.

Simulated and published alleles showed strong mean-composition concordance, with no repeat-unit category significant after multiple-testing correction. PERMANOVA did not detect a centroid shift, whereas PERMDISP indicated greater dispersion among simulated alleles. These results indicate that simulated haplotypes are centered on the published Wenzel composition distribution while spanning a broader per-allele composition range (Figure S9; Table S7).

#### 1.2 SM2: Benchmark design and simulation parameters

##### 1.2.1 Experiment 1: multi-variant, multi-platform benchmark

**Objective:** Benchmarking variant calling tools across variants and sequencing platforms

**Sample design:**

- 220 sample pairs (variant-containing + matched normal control) = 440 samples
- 13 frameshift variants: 100 59dupC pairs (implemented as `dupC`, canonical, ~80% of ADTKD-*MUC1* patients) and 10 pairs each for 12 atypical variants (dupA, insG, insCCCC, insC_pos23, insG_pos58, insG_pos54, insA_pos54, delGCCCA, ins25bp, ins16bp, del18_31, delinsAT)
- VNTR length distribution: Normal(mean=50 repeats per haplotype, SD=17 repeats, clipped to [20, 150]). The core range (35-105 repeats) reflects empirical *MUC1* allele lengths (Vrbacká *et al.*, 2026); extended tails (20-34 and 106-150 repeats, ~15% of samples) stress-test tool performance on extreme lengths.
- **Length distribution rationale.** The Normal(mean = 50, SD = 17) distribution was chosen to span the empirical PacBio full-length *MUC1* allele-length distribution reported by Vrbacká *et al.* (2026), with truncation to the configured simulation bounds. Wenzel *et al.* (2018) informed repeat-unit transition probabilities rather than allele-length sampling. Experiment 2 separately evaluates detection across a wider controlled haplotype-length range.
- Flanking regions: 10 kb upstream and downstream (hg38: chr1:155188487-155192239)
- Random seeds: 5000-5219

Read simulation parameters are summarized in Table S2; tools evaluated in Table S3.

**Read count methodology:** Illumina simulations used a fixed read count of 10000 pairs via the Wessim2-style fragment simulation. Long-read simulations used coverage-based generation (500x amplicon). The MucOneUp software default is 100000 read pairs; all Illumina experiments used 10000 pairs as specified in experiment configuration files.

###### 1.2.2 Experiment 2: VNTR length-asymmetry benchmark

**Objective:** Assess detection sensitivity across haplotype length configurations

**Sample design:**

- 72 59dupC samples (implemented as `dupC`; 12 lengths, 2 conditions, 3 replicates)
- Replicates are independent simulation runs (distinct random seed per pair).
- Haplotype 1: Fixed at 60 repeats
- Haplotype 2: Variable (20, 30, 40, 50, 60, 70, 80, 90, 100, 110, 120, 130 repeats)
- Condition A: Variant on Hap1 (n=36)
- Condition B: Variant on Hap2 (n=36)
- Random seeds: 6000-6071

**Read simulation:** Illumina only, same parameters as Experiment 1

**Analysis focus:** VNtyper 2 Fast mode only (Experiment 1 showed Fast = Normal performance)

###### 1.2.3 Experiment 3: coverage titration benchmark

**Objective:** Assess sensitivity degradation at reduced sequencing depths

**Sample design:**

- Same 220 pairs from Experiment 1

- Illumina platform only, VNtyper 2 Fast mode only
- 4-variant subset: 59dupC (`dupC`), insA_pos54, ins25bp, delGCCCA

**Downsampling method:** BAMs from Experiment 1 were downsampled using `samtools view -s` (SAMtools v1.21).

| Label | Fraction | Effective coverage |
|---|---|---|
| ds75 | 0.75 | ~112.5x |
| ds50 | 0.50 | ~75.0x |
| ds25 | 0.25 | ~37.5x |
| ds12 | 0.125 | ~18.75x |

Effective coverage = fraction multiplied by 150x base coverage.

Coverage titration and runtime/memory analyses are summarized in Figures S4 and S6, respectively.

#### 1.3 SM3: Variant-calling and post-processing protocols

##### 1.3.1 VNtyper 2 mode-specific parameters

All VNtyper 2 analyses used the `vntyper pipeline` subcommand (v2.0.3; archived on Zenodo, concept DOI 10.5281/zenodo.19744166 (Popp and Saei, 2026)). Mode-specific invocations as used in our experiment scripts:

**Fast mode:**

```
vntyper pipeline \
  --bam input.bam \
  -o output/fast_mode/ \
  --reference-assembly hg38 \
  --fast-mode
```

**Normal mode (default):**

```
vntyper pipeline \
  --bam input.bam \
  -o output/normal_mode/ \
  --reference-assembly hg38
```

**Shark mode** (requires paired FASTQ input):

```
vntyper pipeline \
  --fastq1 reads_R1.fastq.gz \
  --fastq2 reads_R2.fastq.gz \
  -o output/shark_mode/ \
  --reference-assembly hg38 \
  --extra-modules shark
```

**adVNTR mode:**

```
vntyper pipeline \
  --bam input.bam \
  -o output/advntr_mode/ \
  --reference-assembly hg38 \
```

```
  --extra-modules advntr
```

##### 1.3.2 Kestrel configuration

VNtyper 2's Kestrel module (Fast, Normal, Shark modes) uses k-mer-based variant detection with internally calibrated parameters (Saei *et al.*, 2023):

- k-mer size: 20 bp
- Minimum k-mer coverage: 3-fold
- Variant confidence threshold: Default (internally calibrated)
- Artifact filtering: Enabled

**Flagged variants:**

1. False Positive 4 bp Insertion (4-base pair insertion artifact resulting from k-mer misalignment at repetition boundaries)
2. Cross mark (a generic artifact indicator for low-confidence assessments)

##### 1.3.3 Classification of filtered calls

Kestrel-flagged VNtyper 2 calls were treated as negative calls in the primary analysis, reflecting VNtyper 2's production filtering behavior. Because the comparator tools do not expose an equivalent artefact filter, Table S1 also reports VNtyper 2 Fast metrics with Kestrel-filter suppression disabled. In this benchmark, the filtered and unfiltered VNtyper 2 Fast rows were identical because no samples carried a Kestrel artefact flag.

###### 1.3.4 Mutation-counter paired enrichment analysis

The Mutation Counter method was previously described as a sequence-counting approach without a reusable software release (Fages *et al.*, 2024); this benchmark used the VNtyper 2 implementation of that method. For each Experiment-1 Illumina BAM, the module streamed aligned read sequences, counted exact target sequences and reverse complements, and emitted variant-specific columns plus the `All` control column. Matched variant-containing and normal samples were compared with one-sided Fisher exact tests for enrichment of the injected variant column against `All` (Virtanen *et al.*, 2020). P-values were evaluated at the pre-specified mutation-counter threshold $P <$ 1.0e-3. The predefined target list supports `dupC`, `dupA`, `insG`, `insCCCC`, and `delinsAT`; pairs carrying unsupported simulated variant classes were retained in the denominator and labelled `unsupported_target`.

###### 1.3.5 WT-vs-WT null analysis

To assess exact-target false enrichment in simulated wild-type samples, we used normal-sample rows from the Mutation Counter count table. Normal samples were randomly paired without self-pairing (seed = 42). For each pair and each supported Mutation Counter target column, we applied the same minimum-count filter as the paired enrichment analysis and computed a two-sided Fisher exact test against the `All` column.

The count-based null analysis produced 0/1100 tests below $P <$ 1.0e-3 (0%), with median P = 1 and mean P = 0.9197 across 5 supported target columns (Figure S8). Because exact target counts in wild-type samples can be zero-inflated, this check estimates false enrichment at the manuscript threshold rather than testing for a uniform P-value distribution.

###### 1.3.6 Matched-coverage re-call analysis

To separate VNTR-length effects from coverage effects, long-allele variant-containing samples from Experiment 2 were downsampled to the median per-base VNTR depth of the short-allele baseline group and re-called with VNtyper 2 Fast. Call agreement before and after downsampling was assessed using Cohen's kappa, and matched-depth Kestrel depth scores were compared with the short-allele baseline using a Mann-Whitney U test.

Matched-depth re-calling results are summarized in Figure S5.

##### 1.4 SM4: Statistical analysis methods

###### 1.4.1 Performance metrics

**Sensitivity (Recall):** TP divided by (TP + FN)

**Specificity:** TN divided by (TN + FP)

**Positive Predictive Value (Precision):** TP divided by (TP + FP)

**Negative Predictive Value:** TN divided by (TN + FN)

**Accuracy:** (TP + TN) divided by (TP + TN + FP + FN)

**F1-Score:** 2(PPV * Sensitivity) / (PPV + Sensitivity)

###### 1.4.2 Correlation analysis

**Pearson correlation coefficient:** Standard implementation

**Significance testing:** Two-tailed *P*-value

**Software:** Python scipy.stats (v1.10+)

**Correlations computed:**

1. Mutation Counter exact-target enrichment and supported-target applicability
2. Hap2 length vs depth score (Experiment 2, Conditions A and B)

##### 1.4.3 Fisher Z comparison

To test whether depth score correlations with Hap2 length differ between Condition A (variant on fixed-length allele) and Condition B (variant on variable-length allele), we applied Fisher's Z-transformation to compare independent Pearson correlations (n=36 per condition, 3 replicates per length). Both conditions showed strong negative correlations between depth score and variable allele length (Condition A: r=-0.693, *P*=2.8635e-6; Condition B: r=-0.625, *P*=4.5418e-5). Fisher's Z-test showed no significant difference between conditions (z=-0.49, *P*=0.6244), indicating that allele length affects detection symmetrically regardless of which haplotype carries the variant.

##### 1.4.4 Ground-truth labels

Ground-truth labels were derived from MucOneUp simulation metadata, which records the applied variant type and repeat position for each simulated sample.

---

### 2 Supplementary Figures

**Figure S1. Cross-platform per-variant sensitivity.** Per-variant detection sensitivity across all tool-platform combinations (n=220 pairs, 13 variants). Heatmap shows sensitivity for each variant type (rows) and tool-platform combination (columns). Short-read tools (VNtyper 2 Fast, Normal, Shark, adVNTR) on Illumina; long-read tools (open-pacmuci on PacBio and ONT, VNTRPipeline on ONT). Green = high sensitivity, red = low. insA_pos54 and delGCCCA are the most challenging variants. VNTRPipeline (ONT) shows the most uniform sensitivity. See Table S1 for aggregate metrics.

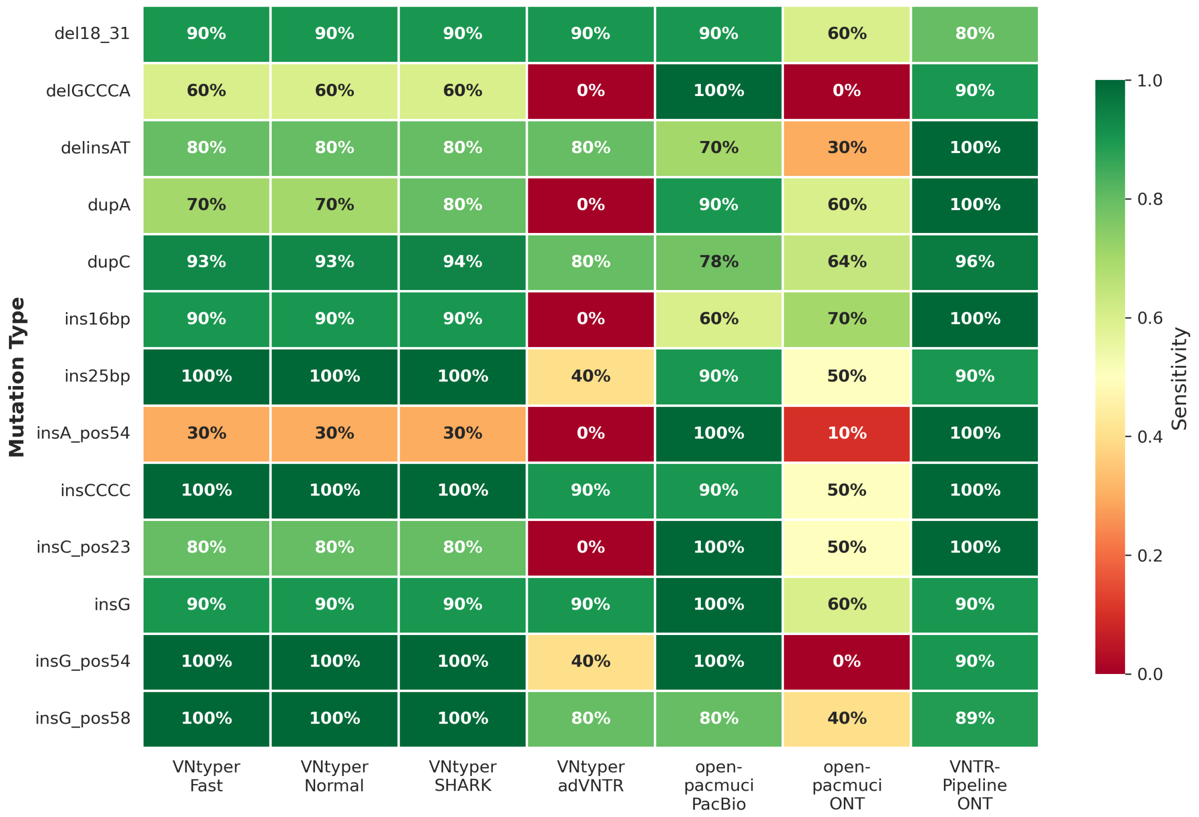

**Figure S2. Mutation Counter paired enrichment analysis.** Fisher's exact tests ([Fages *et al.*, 2024]) on exact Mutation Counter target counts for benchmark pairs covered by the predefined target panel. **A:** Target-specific read counts per evaluated pair (gray = matched normal, blue = variant-containing significant at $P$<1.0e-3, orange = variant-containing not significant), sorted by variant target count. **B:** -$\log_{10}$($P$-value) per evaluated pair (same order; dashed line = $P$=0.001). **C:** Applicability summary showing evaluated supported-target pairs and unsupported-target pairs. Among evaluable pairs, 138/140 (98.6%) showed significant enrichment; 80/220 pairs carried targets outside the Mutation Counter panel.

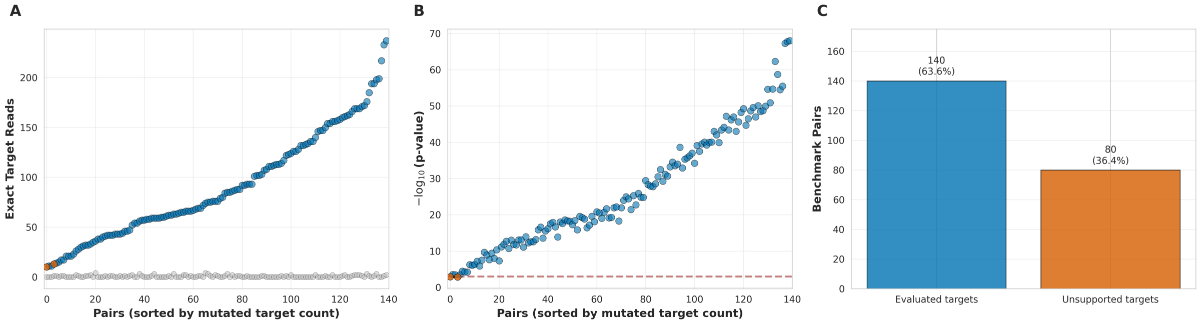

**Figure S3. VNTR length asymmetry effects.** Effect of haplotype length asymmetry on 59dupC detection (n=72, 3 replicates per length, Hap1=60 fixed, Hap2=20-130 variable). **A:** Depth score vs Hap2 length (orange circles = Condition A, variant on Hap1; blue triangles = Condition B, variant on Hap2). Both conditions show strong negative correlations (A: r=-0.693, B: r=-0.625). **B:** Detection rate vs Hap2 length. Overall 94% detection (68/72), with missed calls at extreme lengths. Fisher's Z: no difference between conditions (z=-0.49, *P*=0.6244).

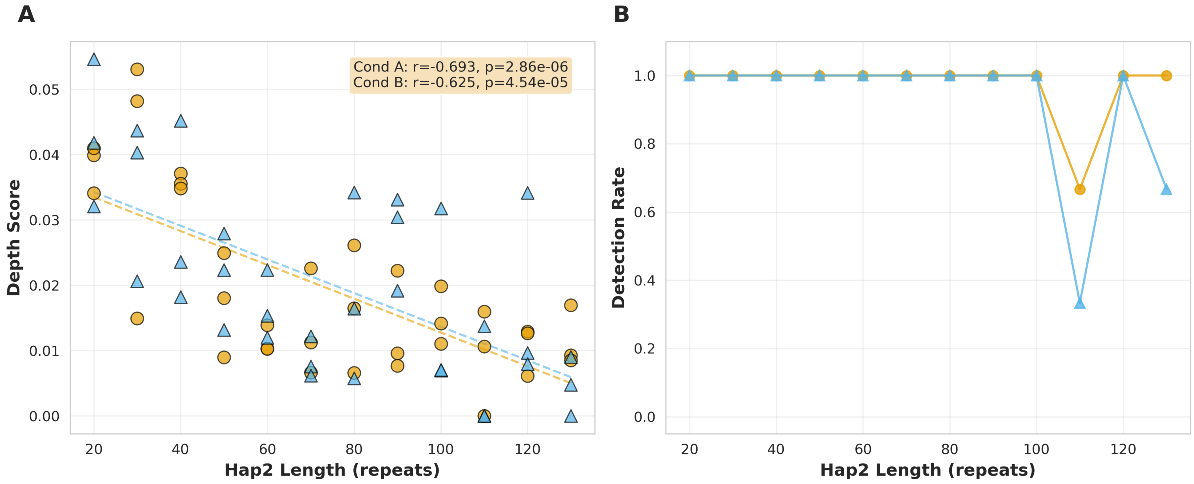

**Figure S4. Coverage titration.** VNtyper 2 Fast sensitivity on Illumina reads downsampled to 4 coverage fractions (220 pairs). **A:** Overall sensitivity vs effective coverage (150x base at 75%, 50%, 25%, 12.5%). Sensitivity declines from 90% at 112.5x to 60% at 18.8x. Shaded band = 95% Wilson CI. **B:** Per-variant curves for 59dupC, insA_pos54, ins25bp, delGCCCA with 95% CI error bars. 59dupC shows steepest decline (94% to 62%); insA_pos54 remains low across all levels.

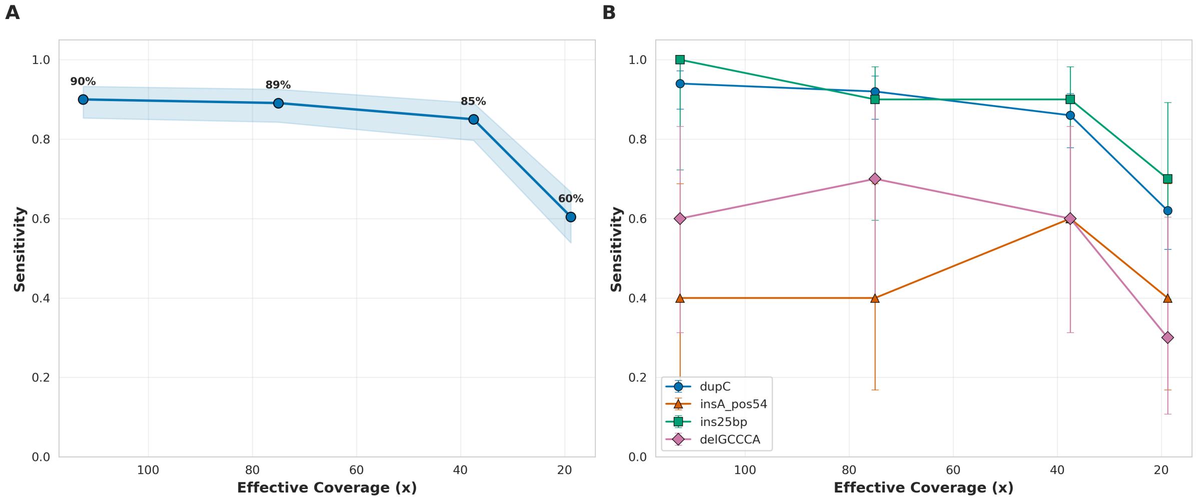

**Figure S5. Matched-coverage re-call for long-allele Experiment-2 samples.** Left: Kestrel depth score versus haplotype-2 repeat count before and after downsampling long-allele samples to the short-allele baseline depth. Right: Kestrel depth-score distributions for the short-allele baseline and matched-depth long-allele calls. Depth scores remained lower for long alleles at matched coverage.

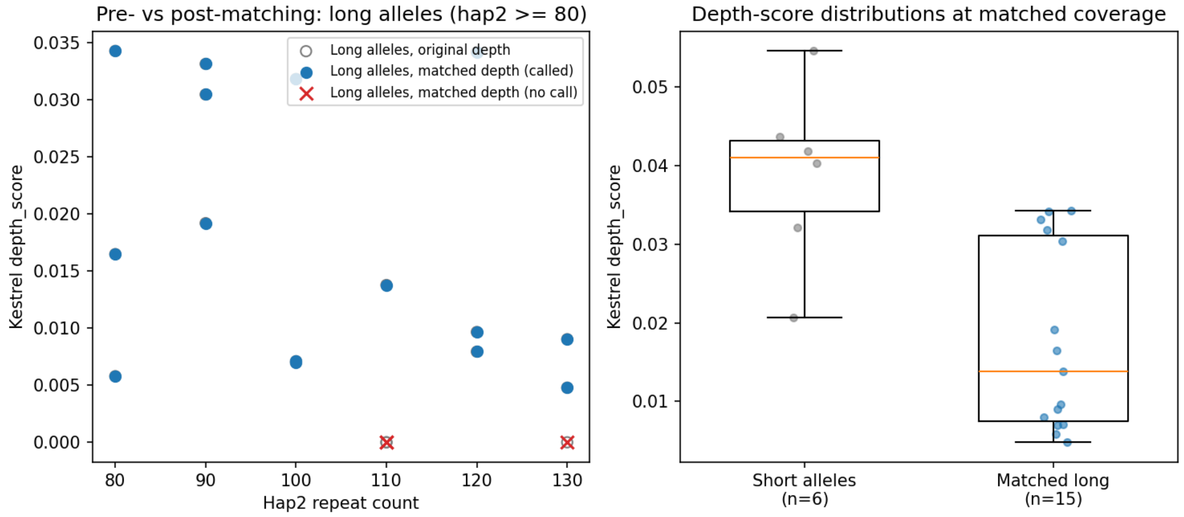

**Figure S 1**

**Figure S6. Runtime and memory.** Processing time and peak memory across tool-platform combinations (n=220 pairs, experiment 1). **A:** Wall time per sample (seconds). **B:** Peak resident set size (RSS) per sample (MB). Box plots with overlaid individual data points. VNtyper 2 adVNTR shows highest runtime variability; long-read tools show moderate resource usage.

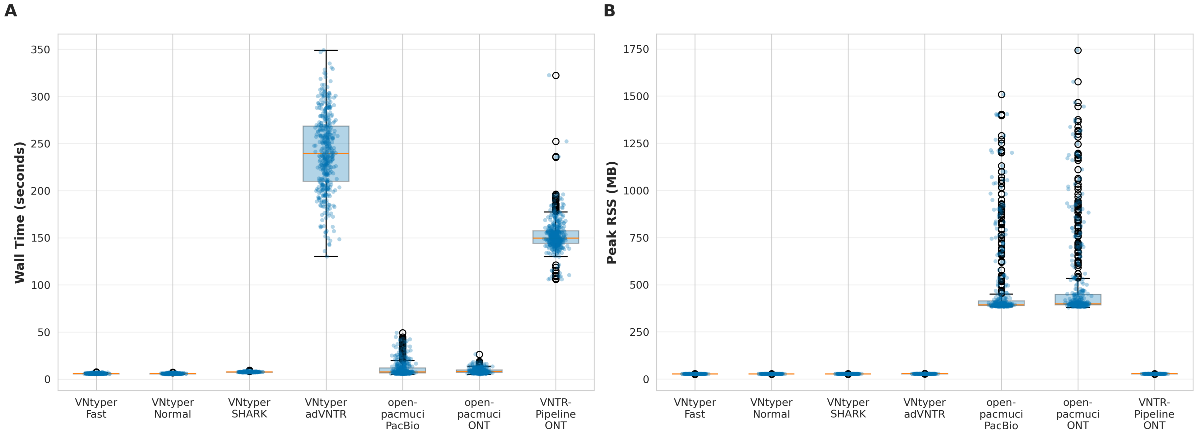

**Figure S7. Composite-score weight-sensitivity analysis.** Classification agreement across repeat-score weights and pathogenicity thresholds. The default configuration uses repeat weight 0.6, composition weight 0.4, and threshold 0.5. Agreement remained high across most tested settings, indicating that binary classification was insensitive to moderate parameter changes. Table S6 gives the numerical grid.

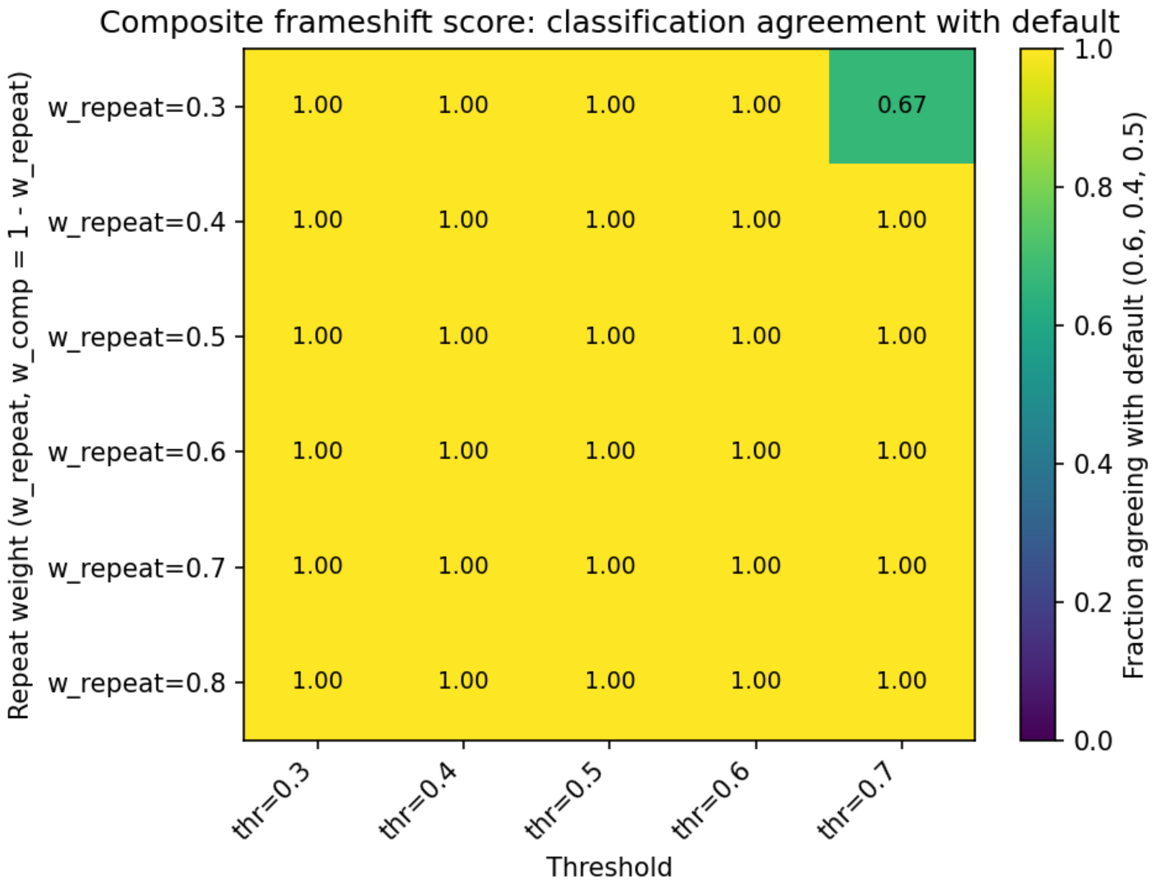

**Figure S8. WT-vs-WT exact-target null analysis.** Two-sided Fisher exact P-values from random pairings of normal Experiment-1 Mutation Counter count rows across 5 supported target columns. The null analysis observed 0/1100 tests below $P < 1.0e\text{-}3$ (0%).

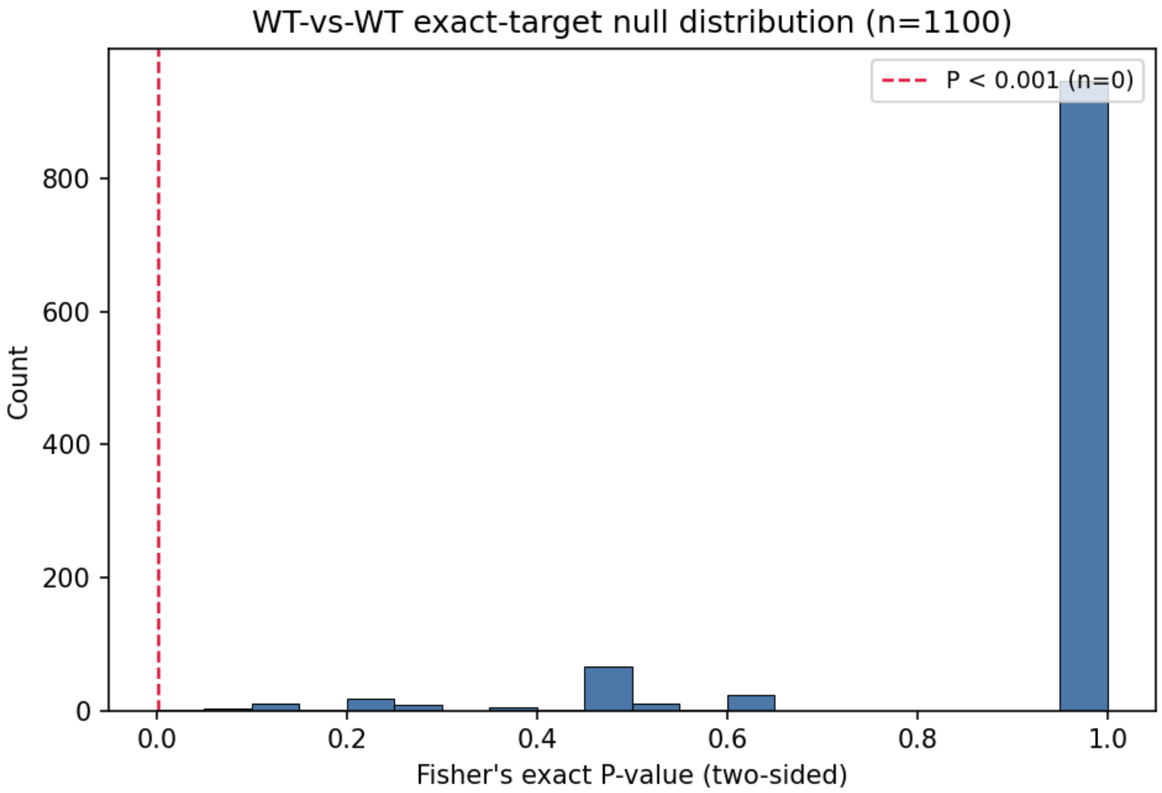

**Figure S9. Per-allele VNTR composition in principal-coordinate space.** Principal-coordinate analysis of Bray-Curtis dissimilarities between Wenzel *et al.* (2018) alleles and MucOneUp-simulated normal haplotypes represented as 28-category repeat-unit composition vectors. Large X markers show group centroids. The groups showed similar centroids and greater dispersion among simulated haplotypes.

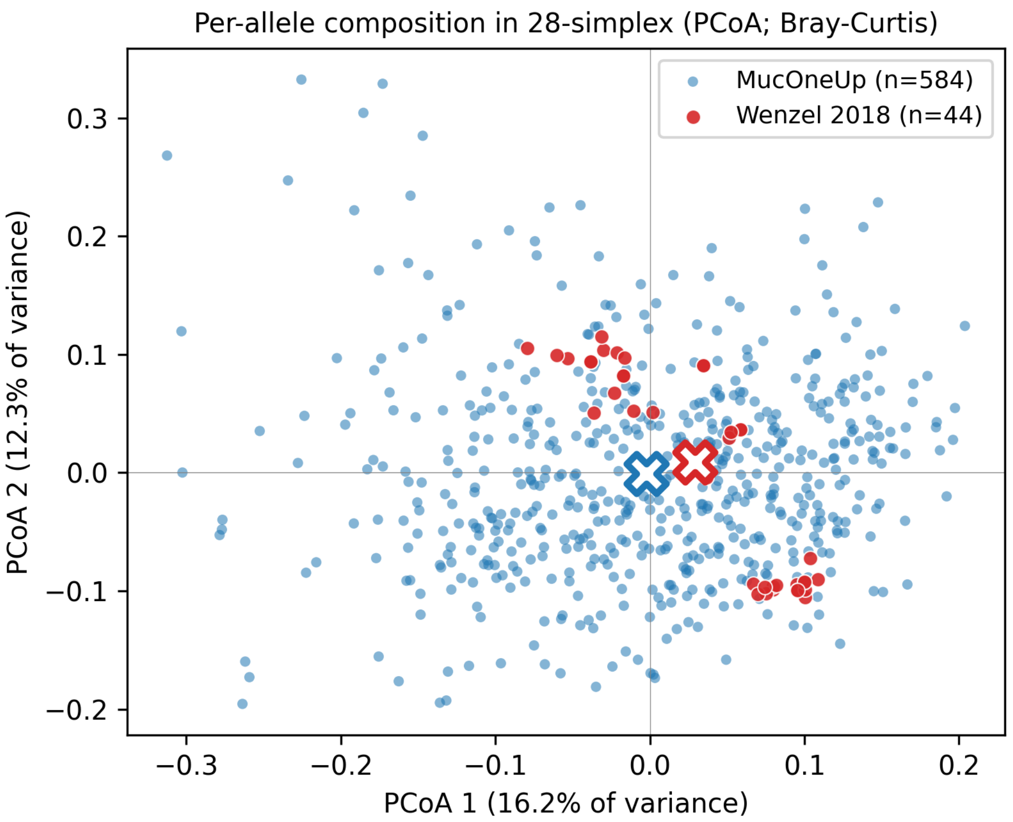

### 3 Supplementary Tables

**Table S1. Performance metrics** for *MUC1* VNTR variant calling across 13 variants (220 pairs, 440 samples). Illumina data were simulated at 150x, and long-read data were simulated as 500x amplicon reads. Normal mode is omitted (identical to Fast). VNTRPipeline reports LOF-only. CI denotes 95% Wilson confidence interval for sensitivity. The VNtyper 2 Fast unfiltered row reports metrics with Kestrel-filter suppression disabled; in this benchmark, filtered and unfiltered VNtyper 2 Fast metrics were identical because no samples carried a Kestrel artefact flag.

| Tool (Platform) | TP | FN | FP | Sens. | Spec. | PPV | F1 | 95% CI |
|---|---|---|---|---|---|---|---|---|
| VNtyper Fast (Illumina) | 192 | 28 | 0 | 0.8727 | 1 | 1 | 0.932 | [0.8222, 0.9105] |
| VNtyper Fast (Illumina, unfiltered) | 192 | 28 | 0 | 0.8727 | 1 | 1 | 0.932 | [0.8222, 0.9105] |
| VNtyper Shark (Illumina) | 194 | 26 | 0 | 0.8818 | 1 | 1 | 0.9372 | [0.8325, 0.9181] |
| VNtyper adVNTR (Illumina) | 131 | 89 | 0 | 0.5955 | 1 | 1 | 0.7464 | [0.5295, 0.6581] |
| open-pacmuci (PacBio) | 185 | 35 | 12 | 0.8409 | 0.9455 | 0.9391 | 0.8873 | |
| open-pacmuci (ONT) | 112 | 108 | 0 | 0.5091 | 1 | 1 | 0.6747 | |
| VNTRPipeline (ONT) | 207 | 11 | 2 | 0.9495 | 0.9908 | 0.9904 | 0.9696 | |

**Table S2. Read simulation parameters.** Three sequencing platforms used for benchmarking (experiment 1). Coverage denotes target depth; alignment tools used for BAM generation.

| Platform | Simulator | Coverage | Alignment |
|---|---|---|---|
| Illumina (Hs-Nova-TruSeq.reseq) | Wessim2-style + ReSeq2 v2.0 (PE) | 150x | BWA-MEM v0.7.17 |
| PacBio HiFi (amplicon) | PBSIM3 v3.0.1 + CCS v6.4.0[a] | 500x | minimap2 v2.24 (map-hifi) |
| Oxford Nanopore (amplicon) | PBSIM3 v3.0.1[b] | 500x | minimap2 v2.24 (map-ont) |

[a] PacBio HiFi: ERRHMM-SEQUEL error model; CCS consensus with 10 passes, minimum 3 passes, minimum read quality 0.99. [b] ONT amplicon: QSHMM-ONT-HQ error model (accuracy mean 0.95). Both amplicon pipelines simulate haplotypes independently with PCR-bias-weighted coverage and unique seeds per allele.

**Table S3. Tools evaluated.** Seven tool-platform combinations benchmarked in experiment 1. VNtyper 2 (Popp and Saei, 2026) extends the original VNtyper (Saei *et al.*, 2023) with Fast and Normal modes (Normal includes unmapped read extraction), integrates code-adVNTR (Bakhtiari *et al.*, 2018; Park *et al.*, 2022) as a detection mode, and implements the SHARK filtering approach (Bensouna *et al.*, 2024) and Mutation Counter method (Fages *et al.*, 2024) as software pipelines. open-pacmuci reimplements the PacBio workflow described by Vrbacká *et al.* (2026); VNTRPipeline is described in (Madritsch *et al.*, 2025).

| Tool (Version) | Platform | Detection Method |
|---|---|---|
| VNtyper 2 Fast (2.0.3) | Illumina | Kestrel k-mer genotyping |
| VNtyper 2 Normal (2.0.3) | Illumina | Kestrel (full pipeline) |
| VNtyper 2 Shark (2.0.3) | Illumina | Kestrel + SHARK filtering |
| VNtyper 2 adVNTR (2.0.3) | Illumina | adVNTR HMM-based |
| open-pacmuci (0.8.0) | PacBio | Clair3 (built-in HiFi model) + allele classification |
| open-pacmuci (0.8.0) | ONT | Clair3 (built-in ONT model) + allele classification |
| VNTRPipeline (2.0) | ONT | Canu assembly + LOF detection |

**Table S4. Two reading frames** following *MUC1* VNTR frameshift variants. The pathogenic frame generates a truncated protein (~85 amino acids downstream of the frameshift) that is trapped in TMED9-containing vesicles of the early secretory pathway (Dvela-Levitt *et al.*, 2019); the non-pathogenic frame terminates later (~189 amino acids downstream). Both distances refer to the position downstream of the frameshift within the repeat unit. Because repeat unit X contains no stop codons in any reading frame, the premature stop codons only appear at the transition from the repetitive VNTR to the terminal block, producing similar truncation regardless of variant position within the array. Values were derived from the ORF interpretation used for the systematic frameshift analysis in Table S5.

| Variant | HGVS Notation | Frame | Stop (AA downstream) |
|---|---|---|---|
| 2 bp del | p.Ala144Profs*85 | Pathogenic | ~85 |

| Variant | HGVS Notation | Frame | Stop (AA downstream) |
|---|---|---|---|
| 5 bp del | p.Ala144Argfs*84 | Pathogenic | ~84 |
| 8 bp del | p.Ala144Cysfs*83 | Pathogenic | ~83 |
| 11 bp del | p.Ala144Hisfs*82 | Pathogenic | ~82 |
| 1 bp del | p.Ala144Profs*189 | Non-pathogenic | ~189 |
| 4 bp del | p.Ala144Thrfs*188 | Non-pathogenic | ~188 |
| 7 bp del | p.Ala144Valfs*187 | Non-pathogenic | ~187 |
| 10 bp del | p.Ala144Serfs*186 | Non-pathogenic | ~186 |

**Table S5. Systematic frameshift analysis** computed using MucOneUp `analyze orfs`. Thirty variants (insertions of 1–15 cytosines and deletions of 1–15 bp) at position 60 of repeat unit X, with 60 VNTR repeats (seed 9999). `ins1C` is the synthetic one-cytosine insertion corresponding to the canonical 59dupC event in this series. WT reference ORF: 1298 AA (frame 2). $\Delta$AA = WT minus variant ORF length. Score = MucOneUp composite frameshift score; values at or above threshold 0.5 are flagged as the predicted pathogenic frame. Frameshifts scoring below threshold produce near-full-length ORFs (small $\Delta$AA) because repeat unit X contains no stop codons in any reading frame; the shifted frame continues through the repetitive region and encounters a stop codon only near the terminal block, yielding a protein of similar length but altered composition that the detector correctly classifies as below threshold. *In-frame control (multiple of 3 bp).

| Variant | Size | Shift | Variant ORF (AA) | $\Delta$AA | Score | Predicted pathogenic frame |
|---|---|---|---|---|---|---|
| ins1C | 1 | +1 | 1210 | 88 | 0.82 | Yes |

| Variant | Size | Shift | Variant ORF (AA) | ΔAA | Score | Predicted pathogenic frame |
|---|---|---|---|---|---|---|
| ins2C | 2 | +2 | 1288 | 10 | 0.15 | No |
| ins3C* | 3 | 0 | 1299 | -1 | 0.10 | No |
| ins4C | 4 | +1 | 1211 | 87 | 0.82 | Yes |
| ins5C | 5 | +2 | 1289 | 9 | 0.15 | No |
| ins6C* | 6 | 0 | 1300 | -2 | 0.10 | No |
| ins7C | 7 | +1 | 1212 | 86 | 0.82 | Yes |
| ins8C | 8 | +2 | 1290 | 8 | 0.15 | No |
| ins9C* | 9 | 0 | 1301 | -3 | 0.10 | No |
| ins10C | 10 | +1 | 1213 | 85 | 0.82 | Yes |
| ins11C | 11 | +2 | 1291 | 7 | 0.15 | No |
| ins12C* | 12 | 0 | 1302 | -4 | 0.10 | No |
| ins13C | 13 | +1 | 1214 | 84 | 0.82 | Yes |
| ins14C | 14 | +2 | 1292 | 6 | 0.15 | No |
| ins15C* | 15 | 0 | 1303 | -5 | 0.10 | No |
| del1bp | 1 | -1 | 1287 | 11 | 0.15 | No |
| del2bp | 2 | -2 | 1209 | 89 | 0.82 | Yes |
| del3bp* | 3 | 0 | 1297 | 1 | 0.10 | No |
| del4bp | 4 | -1 | 1286 | 12 | 0.15 | No |
| del5bp | 5 | -2 | 1208 | 90 | 0.82 | Yes |
| del6bp* | 6 | 0 | 1296 | 2 | 0.10 | No |
| del7bp | 7 | -1 | 1285 | 13 | 0.15 | No |
| del8bp | 8 | -2 | 1207 | 91 | 0.82 | Yes |

| Variant | Size | Shift | Variant ORF (AA) | ΔAA | Score | Predicted pathogenic frame |
|---|---|---|---|---|---|---|
| del9bp* | 9 | 0 | 1295 | 3 | 0.10 | No |
| del10bp | 10 | -1 | 1284 | 14 | 0.15 | No |
| del11bp | 11 | -2 | 1206 | 92 | 0.82 | Yes |
| del12bp* | 12 | 0 | 1294 | 4 | 0.10 | No |
| del13bp | 13 | -1 | 1283 | 15 | 0.15 | No |
| del14bp | 14 | -2 | 1205 | 93 | 0.82 | Yes |
| del15bp* | 15 | 0 | 1293 | 5 | 0.10 | No |

**Table S6. Weight-sensitivity agreement grid.** Fraction of frameshift variants receiving the same binary classification as under the default configuration. Rows vary repeat-score weight, with composition weight defined as 1 minus repeat-score weight; columns vary the pathogenicity threshold. The default cell equals 1.00 by construction. See Figure S7.

| | thr=0.3 | thr=0.4 | thr=0.5 | thr=0.6 | thr=0.7 |
|---|---|---|---|---|---|
| w_repeat=0.3 | 1.00 | 1.00 | 1.00 | 1.00 | 0.67 |
| w_repeat=0.4 | 1.00 | 1.00 | 1.00 | 1.00 | 1.00 |
| w_repeat=0.5 | 1.00 | 1.00 | 1.00 | 1.00 | 1.00 |
| w_repeat=0.6 | 1.00 | 1.00 | 1.00 | 1.00 | 1.00 |
| w_repeat=0.7 | 1.00 | 1.00 | 1.00 | 1.00 | 1.00 |
| w_repeat=0.8 | 1.00 | 1.00 | 1.00 | 1.00 | 1.00 |

**Table S7. Simulated versus published *MUC1* VNTR per-allele repeat-unit com-**

**position.** Mean and standard deviation of per-unit percentages across Wenzel *et al.* (2018) alleles and MucOneUp-simulated normal haplotypes. Absolute mean difference is reported in percentage points. Mann-Whitney U q-values were adjusted using the Benjamini-Hochberg procedure across repeat-unit categories.

| Unit | Wenzel mean % | Wenzel SD | Sim mean % | Sim SD | | Δ mean |
|---|---|---|---|---|---|---|
| X | 47.39 | 6.07 | 44.75 | 9.57 | 2.63 | 0.356 |
| A | 11.77 | 4.35 | 13.13 | 6.15 | 1.37 | 0.365 |
| B | 8.17 | 1.73 | 8.30 | 3.28 | 0.14 | 0.951 |
| C | 5.10 | 2.51 | 4.56 | 2.20 | 0.54 | 0.365 |
| V | 2.23 | 1.43 | 3.16 | 3.98 | 0.93 | 0.866 |
| E | 3.09 | 1.62 | 2.45 | 1.96 | 0.64 | 0.191 |
| G | 2.18 | 1.21 | 2.66 | 2.17 | 0.48 | 0.365 |
| D | 2.61 | 1.44 | 2.15 | 1.83 | 0.46 | 0.356 |
| 1 | 1.76 | 0.48 | 1.95 | 0.80 | 0.20 | 0.365 |
| 2 | 1.76 | 0.48 | 1.95 | 0.80 | 0.20 | 0.365 |
| 3 | 1.76 | 0.48 | 1.95 | 0.80 | 0.20 | 0.365 |
| 5 | 1.76 | 0.48 | 1.95 | 0.80 | 0.20 | 0.365 |
| 8 | 1.76 | 0.48 | 1.95 | 0.80 | 0.20 | 0.365 |
| 9 | 1.76 | 0.48 | 1.95 | 0.80 | 0.20 | 0.365 |
| 7 | 1.70 | 0.54 | 1.95 | 0.80 | 0.25 | 0.365 |
| 4 | 1.70 | 0.54 | 1.91 | 0.86 | 0.20 | 0.365 |
| 6p | 1.27 | 1.05 | 0.98 | 1.13 | 0.29 | 0.191 |
| 6 | 0.49 | 0.62 | 0.97 | 1.13 | 0.48 | 0.078 |

| Unit | Wenzel mean % | Wenzel SD | Sim mean % | Sim SD | | Δ mean |
|---|---|---|---|---|---|---|
| F | 0.96 | 1.12 | 0.60 | 1.08 | 0.37 | 0.097 |
| J | 0.25 | 0.60 | 0.25 | 0.66 | 0.00 | 0.951 |
| I | 0.19 | 0.54 | 0.11 | 0.46 | 0.08 | 0.365 |
| H | 0.06 | 0.30 | 0.09 | 0.39 | 0.03 | 0.951 |
| L | 0.09 | 0.42 | 0.03 | 0.22 | 0.06 | 0.684 |
| N | 0.04 | 0.30 | 0.05 | 0.31 | 0.00 | 0.951 |
| R | 0.04 | 0.30 | 0.05 | 0.31 | 0.00 | 0.951 |
| 4p | 0.05 | 0.35 | 0.05 | 0.29 | 0.01 | 0.951 |
| P | 0.03 | 0.19 | 0.04 | 0.25 | 0.01 | 0.951 |
| Q | 0.03 | 0.19 | 0.03 | 0.24 | 0.01 | 0.955 |

### 4 Supplementary Data

**Data S1.** Complete simulation parameters and benchmarking results (Excel workbook). Contains per-pair haplotype configurations (repeat counts, VNTR lengths, GC content, ordered repeat structure with variant-bearing unit annotation), detection results for all tool-platform combinations (3,220 calls), overall performance metrics (sensitivity, specificity, PPV, NPV with Wilson CIs), per-variant sensitivity breakdowns, count-based Mutation Counter results with raw target-count sheets, and frameshift analysis predictions. Available as supplementary material and at the Zenodo data deposit.
